# Pericytes Repair Engineered Defects in the Basement Membrane to Restore Barrier Integrity in an in vitro Model of the Blood-Brain Barrier

**DOI:** 10.1101/2025.06.23.661146

**Authors:** Michelle A. Trempel, Yimei Du, Louis P. Widom, Emily E. Reitz, Alexis M. Feidler, Pelin Kasap, Britta Engelhardt, Thomas R. Gaborski, Harris A. Gelbard, Niccolo Terrando, James L. McGrath

## Abstract

Pericytes play a key role in the brain where they support the blood-brain barrier (BBB). Their loss has been reported in response to systemic inflammation and neurodegenerative disease. We recently demonstrated that iPSC-derived brain pericyte-like cells (BPLCs) and brain microvascular endothelial cell (BMEC)-like cells collaboratively form a nascent, 3D basement membrane when cultured across a nanoporous membrane^1^. Building on this, we aimed to engineer defects in the basement membrane to investigate whether pericytes could facilitate its repair. In BMEC monocultures, we observed that micropore patterns in nanomembranes created discontinuities in laminin, which destabilized barrier function. Remarkably, the addition of pericytes to the basal side of the membrane restored both laminin integrity and barrier function. Our results align with the role of pericytes as support cells for microvasculature and encourage the use of our tissue barrier platform (the µSiM) to model neurological disorders involving pericyte dysfunction and/or disruption of basement membrane.

---

The blood-brain barrier (BBB) maintains the homeostasis of the brain by regulating cellular and molecular transport between blood and brain, maintaining metabolic homeostasis while restricting toxins and pathogens^2, 3^. The capillary wall is formed by brain microvascular endothelial cells (BMECs) with support from the basement membrane (BM) and pericytes^4^. The barrier is maintained by preventing paracellular diffusion with tight junctions between BMECs^3^ and minimizing transcellular transport via reduced pinocytosis in BMECs^5^. Pericyte loss and increased BBB permeability have been seen with aging^6, 7^, brain injury following systemic inflammation^8^, and various neurodegenerative disorders^7, 9^. Because pericytes have no intrinsic barrier properties, while BMECs do^10^, pericytes are clearly barrier-*supporting* cells. Here we sought to demonstrate that this support role could be modeled *in vitro* by engineering barrier-destabilizing defects in an artificial BM and asking if the addition of pericytes could restore barrier stability.

The connection between pericytes, BM maintenance, and central nervous system (CNS) diseases is well illustrated by the pathophysiological effects of APOE4, the strongest genetic risk factor for late-onset Alzheimer’s disease (AD)^11^. The common isoforms of APOE interact with lipoprotein receptor-related protein 1 (LRP1) differently^12^. While the E2 and E3 isoforms effectively bind to LRP1, suppressing the proinflammatory cyclophilin A (CypA)– matrix metalloproteinase-9 (MMP9) pathway in pericytes, APOE4 interacts weakly with LRP1 leading to BM degradation, loss of cell adhesion to the vascular wall, and vascular dysfunction^12, 13^. Microfabrication and induced pluripotent stem cell (iPSC) technologies allow for the creation of isogenic human-relevant *in vitro* models of tissue health and disease^14^. Unlike animal models, these advanced *in vitro* culture platforms enable reductionist studies of disease mechanisms and opportunities for high throughput discovery of therapeutics interventions. Here we use the µSiM barrier modeling platform, which features ultrathin (100 nm) **si**licon nitride nano**m**embranes (SiM)^1, 15^ with iPSC-derived extended endothelial culture brain method microvascular endothelial-like cells (EECM-BMECs)^16^ and brain pericyte-like cells (BPLCs)^17^. We build on our previous use of the µSiM for BBB modeling^1, 15^ where we demonstrated that EECM-BMECs and BPLCs grown on either side of nanomembranes synthesize a multicomponent BM^1^. In this BM, BPLCs reside within an abluminal 3D fibrous matrix that includes fibronectin and collagen IV as seen *in vivo*^18^, and BPLCs and EECM-BMECs both contribute to a laminin-rich substrate on the abluminal side of the barrier-forming EECM-BMEC monolayer. Our current study leverages the modularity of the µSiM platform to engineer defects of varying size and number in the laminin component of the BM, enabling us to assess whether BPLCs can contribute to repair. We achieve this with customized dual-scale (DS) nanomembranes containing both patterned micropores and self-assembled nanopores^19^. The nanomembranes are thin enough (100 nm)^20^ to enable cell-cell and cell-matrix contact across them and are invisible in light microscopy^1, 21^. Our results show that EECM-BMEC barriers become increasingly destabilized as substrate defects increase in size and number, with 5 µm pores also prompting transmembrane migration. Both barrier destabilization and transmigration are mitigated by the addition of BPLCs, which lead to fibrous laminin expression, restoring BM continuity. The addition of pericytes to DS membranes with a modest number of defects dampens barrier breakdown in response to inflammatory stimulation compared to nanoporous-only (NPN) membranes or membranes with large sized defects. This suggests that direct contact between EECM-BMECs and BPLC-produced matrix is stabilizing against an inflammatory challenge, but that there are limits to the ability of pericytes to provide this stabilization.

Before examining the interplay between pericytes, BMECs and their jointly synthesized BM, we first reproduced our results showing that isogenic EECM-BMECs and BPLCs create a rich 3D matrix when cultured on either side of the µSiM’s ultrathin silicon nitride membrane for 7 days. With nanoscale thinness (100 nm) and high porosity (∼15%) the µSiM membranes do not hinder the exchange of small solutes between apical and basal compartments^21, 22^ and they are optically transparent in light microscopy. Our results confirm the formation of a stratified 3D BM-like structure consisting of sheet-like laminin beneath EECM-BMECs and fibrous fibronectin and collagen IV in the abluminal domain occupied by BPLCs (Figure 1A).

**Figure 1.**
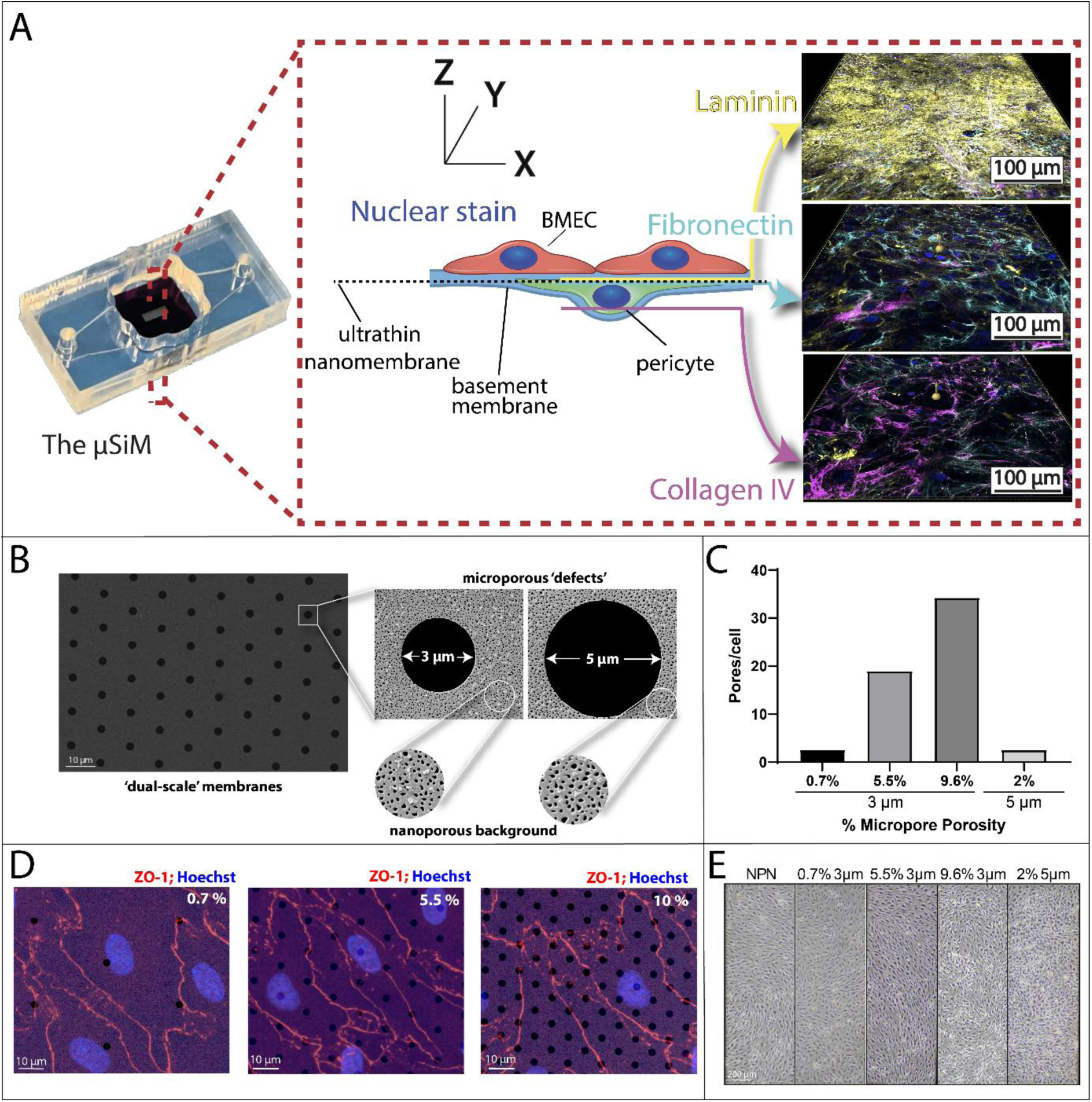
Brain microvascular endothelial-like cells (BMECs) and Brain Pericyte-Like Cells (BPLCs) form a complex 3D basement membrane (BM) on the μSiM which we engineer ‘dual-scale’ (DS) membranes to disrupt. (A) Left – Image of the μSiM device, which features a top well compartment and bottom channel separated by a nanomembrane. Middle – Schematic illustrating the relative placement of the EECM-BMECs (red) and BPLCs (green) on opposite sides of the nanomembrane (dashed line), with colored lines to show the relative position of the BM proteins.

We hypothesized that patterning micropores into nanoporous membranes (Figure 1B) would create a compromised BM and destabilize BBB barrier function. Supporting this approach and hypothesis, we previously developed methods to pattern micropores into freestanding nanomembranes and demonstrated that substrates featuring a high density of micropores do not support endothelial monolayer adhesion under physiological fluid shear stress^19^. Our approach allows us to maintain the high porosity background of nanopores, while modulating the number of micropore substrate defects per cell (Figure 1C, D). Importantly, none of the micropore patterns impacted the ability to grow confluent EECM-BMEC monolayers over 6 days in static conditions (Figure 1E).

Right – Confocal microscopy images of EECM-BMEC-like cells cocultured with BPLCs and stained for the BM proteins laminin (yellow), fibronectin (cyan), and type IV collagen (magenta), along with nuclear stain Hoechst (blue). The images, acquired across the nanoporous membrane region of the chip (as shown in the middle diagram), reveal the formation of three distinct layers: a laminin sheet beneath the EECM-BMEC-like cells, a fibronectin-rich matrix embedding the BPLCs, and collagen IV beneath and through the BPLCs. (B) Scanning electron microscopy (SEM) imaging of a ‘dual-scale’ (DS) membrane, with cutaways highlighting the relative sizes of the micropores and the nanoporous background. (C) Quantification of micropore density per cell (using an average cell area of 2,313 μm^2^ based on averaging cell areas from junctional staining) of for each of the DS membrane conditions that will be used, including their pore size and percent microporosity. (D) SEM images of DS membranes with 3 μm micropores at varying densities (0.7% to 9.6% microporosity), overlaid with confocal microscopy images of EECM-BMECs, fluorescently stained for zona occludins-1 (ZO-1) and nuclei on NPN membranes. The overlay of the fluorescent imaging demonstrates the increasing number of micropores per cell as microporosity increases. (E) Phase-contrast microscopy images of EECM-BMECs grown on membranes with different density and sizes of micropores, showing that the cells can form a confluent monolayer on all of the membrane types.

Immunofluorescence analysis revealed discontinuities in the laminin deposited by EECM-BMECs that correspond to the micropore patterns of the engineered membrane (Figure 2A, B). Quantitative analysis further confirmed this by showing that the distance between voids in the laminin coverage corresponded to the designed spacing of membrane micropores (Figure S1). The impact of patterned defects on overall laminin coverage can be quantified as the number of pixels in an image displaying a positive laminin signal (see Methods). When normalized to NPN membranes, the laminin coverage remained above 85% except for high density (9.6%) 3 μm membranes and 5 μm membranes (73% and 60% respectively) (Figure 2C). Notably, despite the number of micropores being roughly the same between the 5 µm membranes and the lowest density 3 μm membranes, the 5μm membranes cause a greater deficit in laminin coverage. This result indicates that both a high number of small pores and a small number of large pores can create a deficit in laminin coverage.

**Figure 2.**
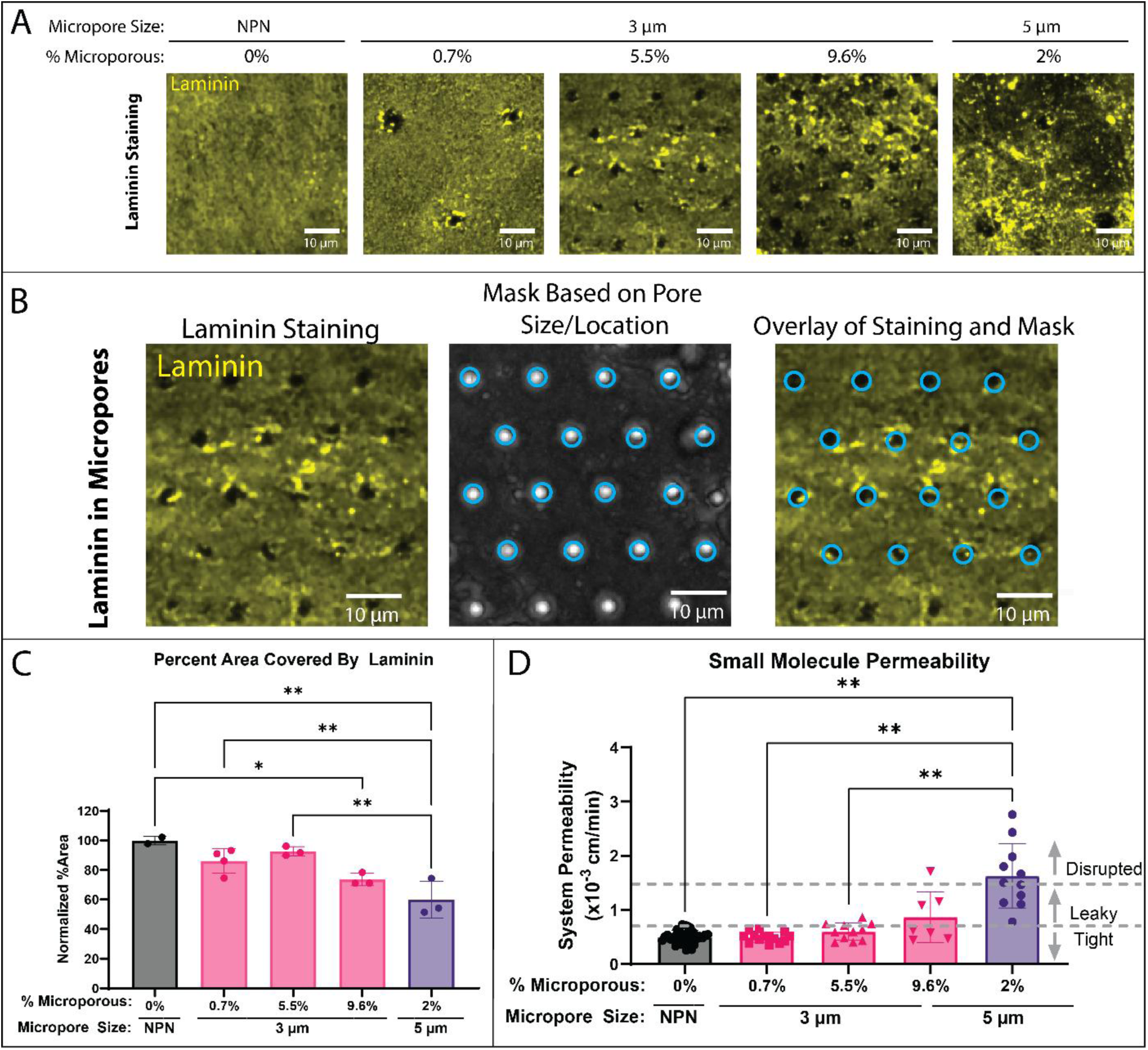
Micropores create corresponding defects in the laminin layer of the BM, correlating with barrier function. (A) EECM-BMECs were cultured on μSiM devices for 6 days and then stained for laminin (yellow). Confocal microscopy was used to acquire images and representative z-maximum intensity projections are shown here (scale bar = 10 μm). Membranes with increasing density or size of micropores show increasing defects in the laminin layer. (B) Overlay of laminin staining with a mask of the pores based on a differential interference contrast (DIC) image of pores reveals that the laminin defects align precisely with the location of the micropores. (C) Quantification of laminin coverage was performed by measuring the percent area with 75% or greater of the mean fluorescent intensity for each image, which was then normalized to the percent area covered in the NPN images. Ordinary one-way ANOVA was used for statistical analysis. (D) Barrier permeability to lucifer yellow small molecule dye (457 Da) over one hour was measured on membranes of different pore sizes and densities and classified as ‘tight’, ‘leaky’, or ‘disrupted’ based on previous work^1^. Brown–Forsythe and Welch ANOVA tests were used for statistical analysis. Statistics: * p≤0.05, ** p≤0.01.

We next asked if the lack of continuity in the laminin surface might induce barrier destabilization. Using our established small molecular permeability assay with lucifer yellow (457 Da)^21, 23^ we found that barrier permeability increased from ‘tight’ to ‘leaky’ as the density of 3 μm pores increased, with barriers cultured on 5 μm pore membranes most consistently measuring in the ‘leaky’ or ‘disrupted’ categories (Figure 2D). Interestingly, at 0.7% porosity with 3 µm pores, all the measurements fell in the ‘tight’ category, consistent with NPN membranes. This suggests EECM-BMEC are able to overcome a very low density of small micropores. Overall, our correlative study appears to support the hypothesis that engineered membrane defects destabilize EECM-BMEC barrier function by inducing discontinuities in BM coverage.

Given the role of pericytes as vascular barrier support cells, and our prior demonstration that BPLCs contribute significantly to a nascent BM formed in coculture with EECM-BMECs on the µSiM platform, we next asked if the addition of BPLCs could ‘rescue’ the barrier function of EECM-BMEC challenged by micropatterned substrate defects. We found that micropore-patterned defects in laminin were far less obvious in immunofluorescence co-culture images (Figure 3A) and that the nature of the laminin matrix changed from sheet-like to fibrous with the addition of micropores. Interestingly, fibrous laminin was not seen in BPLC monocultures, with co-cultures on NPN, or on confocal slices taken far from the EECM-BMEC monolayer (Figure S2). Quantifying laminin immunofluorescence images in the micropores shows fluorescence recovery on each type of membrane studied (Figure 3B). Using differential interference contrast (DIC) imaging to identify micropore locations in monocultures and cocultures (see Figure 2B) we created a micropore-specific mask and quantitatively confirmed that the presence of BPLCs eliminated laminin defects in every case. The percent area metric used above did not show significant laminin recovery except in the case of 5 µm pores (Figure 3C). Returning to our metric of barrier function, small molecule permeability also showed a return to baseline levels in every instance with the addition of BPLCs (Figure 3D).

**Figure 3.**
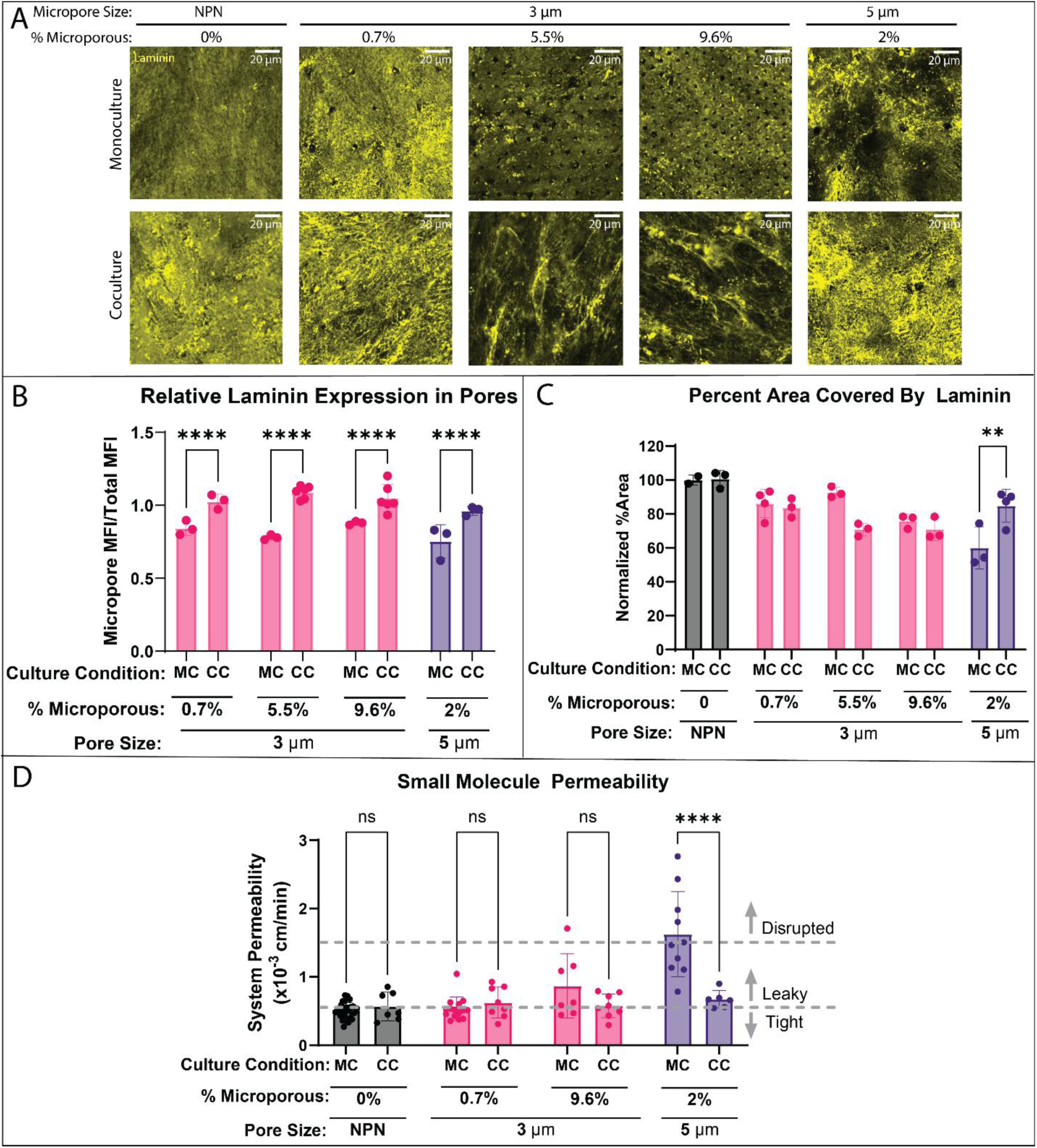
Pericyte addition to DS devices restores laminin coverage and barrier function. (A) Confocal imaging of immunofluorescently stained laminin (yellow) in both BMEC monocultures (MC) and EECM-BMEC/BPLC cocultures (CC) on membranes with increasing densities and sizes of micropores. Scale bar = 20 μm. (B) Quantification of the relative laminin expression in the micropores compared to the total image. Using DIC imaging to identify micropore regions of interest, the average of the mean fluorescent intensity (MFI) of laminin in the micropores was divided by the MFI of the whole image to calculate a ratio of laminin expression in the micropores relative to the total image. Statistical analysis was performed with a 2-way ANOVA test. (C) Quantification of laminin coverage, defined as the percent area with fluorescence intensity greater than 75% of the MFI, normalized to the percent area of MC NPN, shows the addition of BPLCs significantly restores that coverage on the 5 μm DS membranes. Statistical analysis was performed with a 2-way ANOVA test. (D) Small molecule permeability, measured using lucifer yellow (457 Da), reveals a statistically significant increase in permeability in the 5 μm MC condition which is rescued by the addition of BPLCs. Statistical analysis was performed with a 2-way ANOVA test. Statistics: * p≤0.05, ** p≤0.01, *** p≤0.001, **** p≤0.0001.

Our data suggest that micropore size and density can destabilize EECM-BMEC barriers through different mechanisms. This follows from the fact that 5 µm pores only add 2% porosity to nanomembranes (compared to up to 10% for 3 µm pores) and yet they create the greatest deficits in laminin coverage and the most consistently destabilized barriers. Microscopy of EECM-BMEC cells on 5 μm DS membranes revealed an unusual pattern of crisscrossed endothelial junctions and stacked nuclei that were not seen in NPN membranes or 3 μm DS membranes (Figure 4A). This indicated that EECM-BMECs were capable of transmigrating through 5 μm pores to form a monolayer on both sides of the membrane. It is interesting that transmigration resulted in a higher permeability rather than a double endothelial barrier with lower permeability. The addition of BPLCs to create a coculture eliminated transmigration, possibly explaining the restoration of baseline barrier function and recovery of laminin coverage in the case of 5 µm DS membranes (Figure 4B).

**Figure 4.**
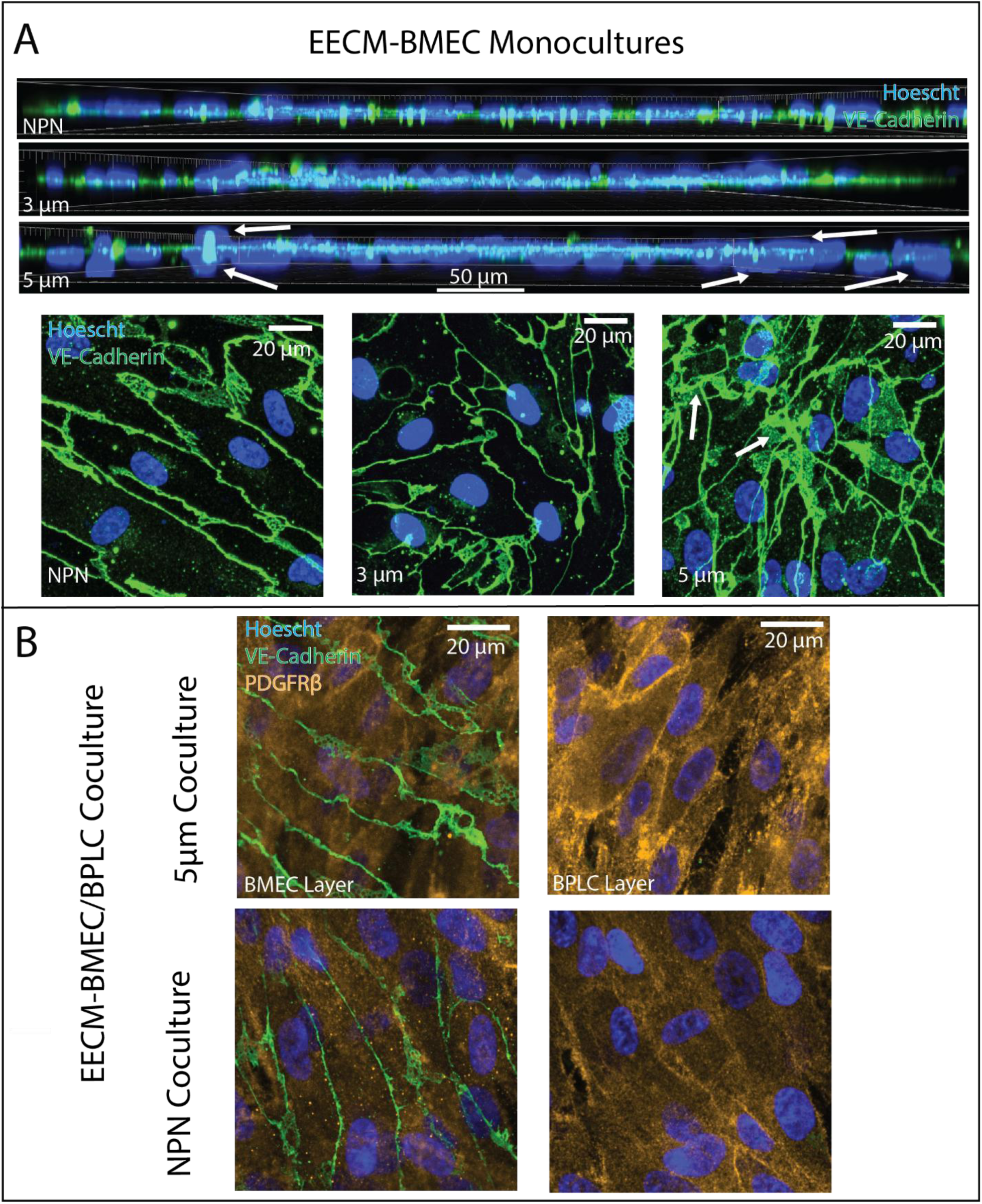
5 μm pores permit BMEC migration through the membrane, which is mitigated by the addition of BPLCs. (A) Confocal imaging of immunofluorescently stained Vascular Endothelial Cadherin (VE-Cadherin) and Hoechst (nuclear stain) on 3 different membrane conditions, NPN, low porosity 3 μm DS, and 5 μm DS. Both side views and top views indicate that while BMECs form a single monolayer in the NPN and 3 μm DS devices, on the 5 μm DS devices they show overlapping nuclei and crisscrossing junctions, suggesting multiple layers of BMECs. (B) Confocal imaging of immunofluorescently stained of VE-Cadherin (used here as a BMEC marker), platelet derived growth factor β (PDGFRβ, a BPLC marker), and Hoechst (nuclear stain), of the BMEC and BPLC layers on NPN and 5 μm DS devices. These images show that these layers are clearly segregated, with BMECs forming a single monolayer which remains above the BPLCs in the CC devices, suggesting that the addition of BPLCs prevents BMEC transmigration.

**Figure 5.**
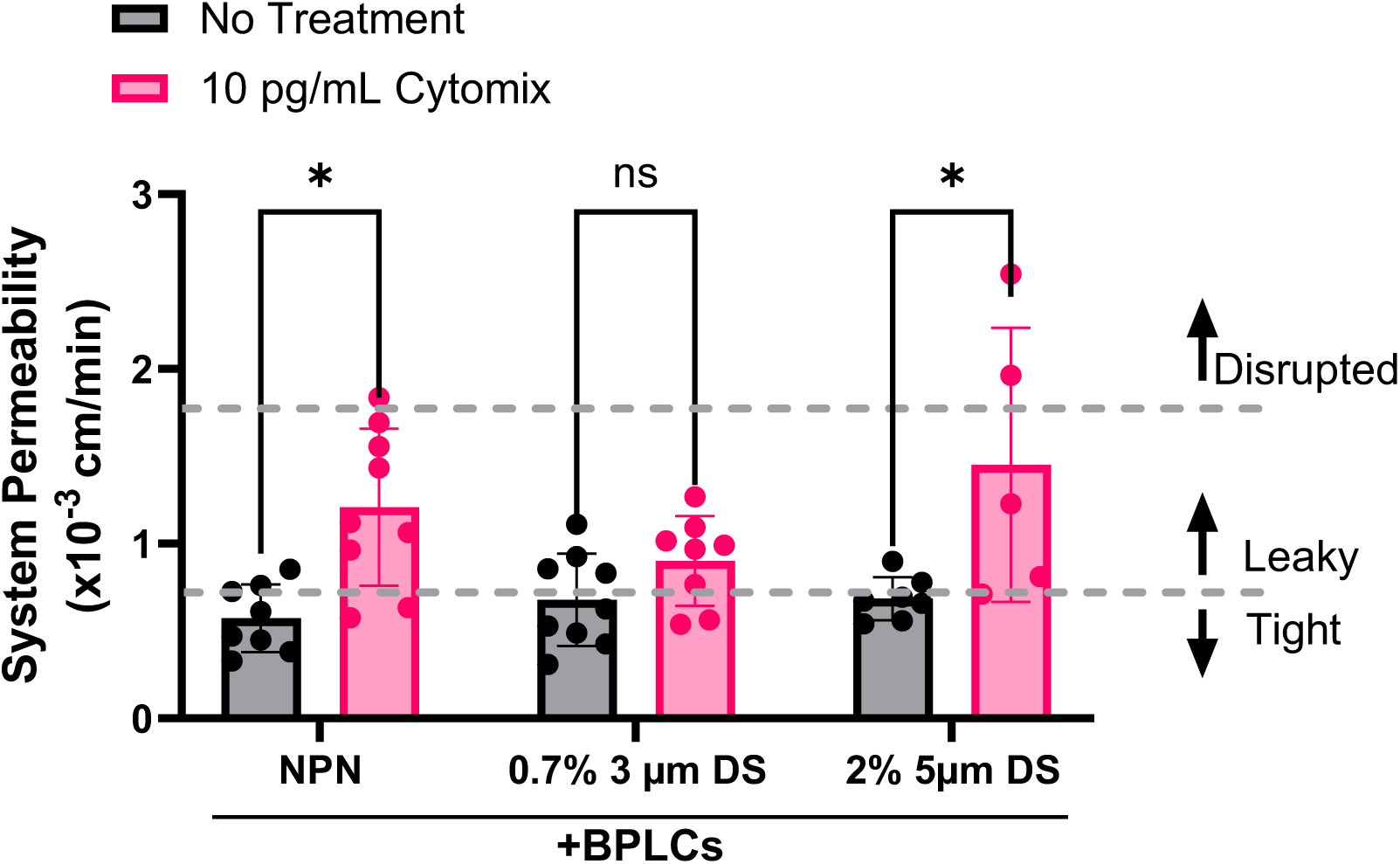
Pro-inflammatory cytokine treatment increases permeability in coculture devices, which is mitigated somewhat by 3 μm pores, but not by 5 μm pores. Small molecule permeability assay (lucifer yellow) results of coculture devices with three types of membranes NPN, 0.7% porosity 3 μm DS and 2% porosity 5 μm DS. Treatment with cytomix (an equimolar mix of TNFα, IL-1β, and IFN-γ) at 10 pg/mL, resulted in a statistically significant increase in permeability for the NPN and 5 μm DS devices. However, that increase is not statistically significant on the 3 μm DS devices, suggesting that low porosity 3 μm pores may mitigate barrier disruption under cytokine stimulation. Statistical analysis was performed using a 2-way ANOVA test. Statistics: * p≤0.05, ns = not significant.

Our laboratory is interested in applying the μSiM to understand and to develop therapies aimed at preventing the transduction of systemic inflammation into brain injury in acute disease settings such as sepsis^24^ and post-operative delirium^25^. We previously found that EECM-BMEC/BPLC cocultures on low porosity 3 μm DS membranes exhibited reduced neutrophil transmigration in response to an cytokine stimulation compared to EECM-BMEC monocultures^1^. In this study, we measured permeability changes following a cytokine stimulation (10 pg/mL each of TNFα, IL-1β, and IFN-γ) for 16-20 h, on NPN, low porosity 3 µm DS, and 5 µm DS membranes and found a similar result. Specifically, we see a dampened permeability increase (not reaching significance) in the case of low porosity 3 µm micropores, but not 5 µm micropores. This limited study illustrates that pericyte reinforcement of barrier function may only be visible under stimulation, as achieving baseline ‘tight’ permeability values on low porosity 3 µm membranes does not require BPLCs^1^. A dampened response on microporous membranes compared to NPN suggests that cell-cell or cell-matrix contact across the membrane is part of the mechanism of stabilization. The high permeability for stimulated cocultures on 5 µm micropores may reflect limits in the capacity of pericytes to support barrier integrity when BM defects are substantial.

Given the critical role of pericytes and the BM in maintaining BBB integrity, our model and its derivatives are well-suited for studying BBB destabilization in disease contexts. These platforms provide a powerful tool for identifying molecular targets for therapeutic intervention and screening of compounds that promote barrier restoration or reinforcement. Particularly valuable is our ability to model ‘metastable’ BBB states—conditions that appear intact under basal conditions but become destabilized following pericyte loss or BM compromise during inflammatory challenges. This feature is especially relevant for modeling the pathophysiology of systemic inflammation-induced brain injury as well as potential mechanisms of barrier dysfunction during aging. Individuals most vulnerable to cognitive decline following acute illness or infection often have predisposing conditions such as advanced age or neurodegenerative disease^26^. In these subjects, latent BBB vulnerabilities may only be unmasked under inflammatory stress^27^. By recreating these hidden susceptibilities *in vitro*, these models have the potential to identify neuroprotective strategies that can be applied early during systemic inflammation—prior to the onset of irreversible brain injury.

## Methods

### Assembling the µSiM Device

The dimensions and assembly of the µSiM have been described in detail previously^15, 28^. Briefly, chips supporting membranes were obtained from SimPore Inc.: NPN (NPSN100.C-1LZ.0), 0.7% 3 μm DS (DSSN100.C-1LZ.0-3.0A1), and custom membranes created by SimPore according to^19^. These chips were bonded to component 1 (upper acrylic component, Aline Inc.) using pressure sensitive adhesive then this was bonded to component 2 (lower cyclic olefin component, Aline Inc.). This was performed under sterile conditions within a biosafety cabinet and the assembled device was exposed to UV light for 20 minutes post-assembly to ensure sterility.

### EECM-BMEC Cell Culture

EECM-BMECs were generated from IMR90-4 hiPSCs (WiCell, Madison Wisconsin) as described here^16^. During the extended endothelial cell culture, the cells were expanded in 6-well tissue culture plates pre-coated with collagen IV (100 µg/mL) reconstituted in water. Cells were cultured in hECSR which consists of human endothelial serum-free medium (Gibco), serum-free B-27 supplement (1x, Gibco) and human basic fibroblast growth factor (20 ng/mL). Media changes were performed every 2-3 days and cells were kept at 37 °C with 5% CO_2_ and 95% humidity. Cells were considered EECM-BMECs and used for assays between passages 4 and 7.

### BPLC Cell Culture

BPLCs were also generated from IMR90-4 hiPSCs as described here^17^. The cells were expanded in uncoated 6-well plates and maintained in Essential 6 (E6) medium with 10% fetal bovine serum (FBS). Media changes were performed daily, and cells were used for assays between Day 22 and Day 45 of differentiation.

### Coculture of EECM-BMECs and BPLCs in the µSiM

EECM-BMECs and BPLCs were cocultured as described in detail here^23^. Devices were washed with sterile water using 20 µL in the bottom channel and 100 µL in the top well. The bottom channels were coated with 20 µL of collagen IV solution (800 µg/mL) (Sigma-Aldrich), while the top wells were coated with a 100 µL mixture of collagen IV (400 µg/mL) and bovine fibronectin (100 µg/mL, Gibco) diluted in water. Devices were incubated with coating solutions for at least 2 hours at 37 °C, followed by washing with E6 medium supplemented with 10% FBS. BPLCs were seeded in the bottom channel of devices at a density of 14,000 cells/cm^2^. Immediately after seeding, devices were inverted. After 2 hours of incubation, the media was changed, and devices were flipped upright. The following day, devices were washed with hECSR medium, and BMECs were seeded in the devices at a density of 40,00-45,000 cells/cm^2^. On the 6^th^ day of BMEC culture, devices were used for assays.

### Small Molecule Permeability Assay

On day 6 of BMEC culture in the µSiM, a small molecule permeability assay was conducted using 150 µg/mL Lucifer Yellow (LY), 457 Da (Invitrogen). The media in the top well was replaced with 100 µL of 150 µg/mL LY solution in hESCR and the devices were incubated for 1 hour at 37 °C. Immediately after incubation, the LY solution was removed and a P200 pipette tip containing 50 µL hECSR was placed in one port connected to the bottom channel of the µSiM. Then a second P200 pipette was inserted into the remaining port of the µSiM and 50 µL was collected and transferred into a black 96-well plate. This process was repeated for each µSiM device, and a standard curve of LY dilutions weas also added to the plate. Fluorescence intensity was measured on a plate reader (TECAN, Mannedorf, Switzerland). System permeability, P_S_, was calculated using Equation (1):

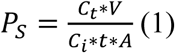

where C_t_ is the concentration of fluorescent small molecule in the bottom channel at time, t, V is the volume transferred to the 96-well plate, C_i_ is the initial concentration of fluorescent small molecule added to the top well, and A is the membrane area.

### VE-cadherin and PDGFRβ Immunofluorescent Staining

Devices were fixed using 4% paraformaldehyde for 15 minutes at room temperature, followed by 3 washes with PBS. Devices were blocked using 5% goat serum + 0.4% Triton X-100 for 30 minutes at room temperature, after which the devices were washed 3 more times with PBS. Primary antibodies – mouse α-human VE-cadherin and rabbit α-human PDGFRβ– were prepared at 1:50 and 1:100 dilutions respectively in blocking solution and added to devices for 1 hour at room temperature. Following incubation devices were washed 3 times with PBS. Secondary antibodies – goat α-Mouse IgG Alexa Fluor 488 and goat α-rabbit IgG Alexa Fluor 568 – were both diluted at 1:200 in blocking solution and incubated on devices for 1 hour at room temperature. After incubation, devices were washed three times using PBS and stained for nuclei using Hoechst 33342 (1:10000 dilution). After Hoechst staining, PBS was added to the top wells and bottom channels of devices and devices were stored at 4 °C until imaging. See Table S1 for antibody details.

### BM Staining

BM proteins were labeled live. Primary antibodies – mouse α-human collagen type IV Alexa Fluor 647; mouse α-human fibronectin, Alexa Fluor 488; and rabbit α-human laminin – were diluted in hESCR with 10% FBS at concentrations of 1:100, 1:200, and 1:100 respectively. Antibody solutions were added to both the top well and bottom channels of devices and devices were incubated for 2 hours at 37 °C, 5% CO_2_ protected from light. Devices were then washed with PBS and fixed with 4% paraformaldehyde for 15 minutes at room temperature. Devices were washed three times with PBS and blocked using 10% goat serum for 10 minutes. Secondary antibody goat α-rabbit Alexa Fluor 568 and Hoechst 33342 were diluted at 1:200 and 1:10,000, respectively, in blocking solution. Devices were incubated in this solution for 1 hour at room temperature. Devices were washed three times in PBS and stored at 4 °C until imaging. See Table S1 for antibody details.

### Imaging Using Confocal Microscopy

Devices were imaged using the Andor Dragonfly Spinning Disc Confocal Microscope in the High Content Imaging Core at the University of Rochester. The confocal microscope stage (Abingdon, UK) was attached to a Nikon TiE microscope (Nikon Corporation, Tokyo, Japan) and a SONA sCMOS (Andor Technology, Belfast, UK) camera. Imaging was performed using the FarRed AF647, DAPI, Green AF488, and Red AF568 fluorescence channels as well as the DIC channel. Z-stack images were captured on 40x magnification using optical slices of 0.2 µm starting below the BPLC layer until the top of the BMEC layer.

### Imaging Analysis

Image analysis was performed using FIJI software^29^. First, maximum intensity z-projections were generated from confocal stacks. The mean fluorescent intensity (MFI) of the entire laminin image was then calculated. A threshold value was determined by calculating 75% of that MFI which was used to identify and quantify the percent area with greater than 75% of the MFI. This was then normalized to the average percent area observed in the NPN monoculture images. Separately, the DIC image of the micropores was processed using the “Analyze Particles” function to generate regions of interest (ROIs) of each pore. Then these ROIs were applied to the laminin image and the MFI within each pore was calculated and averaged. This number was then divided by the total MFI in the laminin image to calculate the ratio of MFI within pores to the total MFI.

## ASSOCIATED CONTENT

### Supporting Information

The follow files are available free of charge.

Table of all antibodies used with relevant conditions and concentration (Table S1); Analysis of laminin defects to demonstrate equivalence to engineered micropores (Section S1); Images of laminin produced by BPLCs alone (Section S2) (PDF).

## AUTHOR INFORMATION

### Author Contributions

The manuscript was written through contributions of all authors. M.A.T., J.L.M, H.A.G., N.T., B.E., and T.R.G. conceptualized and designed the experimental studies. M.A.T. performed the experimental studies with L.P.W. (BPLC monocultures and staining), E.E.R. (some coculture devices and staining), and P.K. (differentiating EECM-BMECs). M.A.T., J.L.M, and A.M.F. designed the image analysis. M.A.T. and Y.D. performed the image analysis. M.A.T. and Y.D. wrote the manuscript draft. All authors contributed to revisions of the manuscript

All authors have given approval to the final version of the manuscript.

## Funding Sources

M.A.T. and H.A.G. were supported by RF1AG079138. L.P.W., T.R.G, A.M.F., P.K., and B.E. were supported by NIH R33 HL154249. E.E.R. was supported by UH3TR003281. N.T. was supported by RF1AG079138 and 2R01AG057525-06A1. J.L.M. was supported by NIH R33 HL154249 and NIH U2CTR004861.

## Notes

The authors declare the following competing financial interests: J.L.M. is a cofounder of SiMPore and holds an equity interest in the company. SiMPore is commercializing ultra-thin silicon-based technologies including the membranes used in this study. B.E. is an inventor on a provisional U.S. patent application (63/185815) related to the methodology of EECM-BMEC-like cell differentiation.

## ACKNOWLEDGMENTS

The authors thank the High Content Image Core (University of Rochester) for help with fluorescent imaging and use of the Dragonfly Spinning Disk Confocal. The authors used chat.rochester.edu for edits of the drafted manuscript.

## Notes

### Competing Interest Statement

The authors declare the following competing financial interests: J.L.M. and T.R.G. are cofounders of SiMPore and holds an equity interest in the company. SiMPore is commercializing ultra-thin silicon-based technologies including the membranes used in this study. B.E. is an inventor on a provisional U.S. patent application (63/185815) related to the methodology of EECM-BMEC-like cell differentiation.

