## Supplemental Table 1, Supplemental Figure 1, Supplemental Figure 2 for "Pericytes Repair Engineered Defects in the Basement Membrane to Restore Barrier Integrity in an in vitro Model of the Blood-Brain Barrier"

**Table S1.** Antibodies used for immunofluorescence staining

| Antibodies | Fixative | Clone | Source | Cat. N. | Dilution |
|---|---|---|---|---|---|
| Mouse Anti-Human VE-Cadherin IgG2B | 4% Paraformaldehyde | 123413 | R&D Systems | MAB9381 | 1:50 |
| Rabbit Anti-Human PDGFRβ IgG | 4% Paraformaldehyde | 28E1 | Cell Signaling Technology | 3169 | 1:100 |
| Mouse Anti-Human Collagen Type IV IgG2b, κ, Alexa Fluor 647 | live | 1042 | Invitrogen | 51-9871-82 | 1:100 |
| Rabbit Anti-Human Fibronectin IgG1, Alexa Fluor 488 | live | FN-3 | Invitrogen | 53-9869-82 | 1:200 |
| Rabbit Anti-Human Laminin IgG | live | Polyclonal | Invitrogen | PA1-16730 | 1:100 |
| Goat Anti-Mouse IgG Alexa Fluor 488 | N/A | N/A | Invitrogen | A11001 | 1:200 |
| Goat Anti-Rabbit IgG Alexa Fluor 568 | N/A | N/A | Invitrogen | A11011 | 1:200 |

Supplementary Section S1: Validation of the distance between laminin defects.

To more thoroughly demonstrate the micropores are causing the defects seen in the monoculture devices, beyond simply aligning defects with images of the micropores, an analysis was performed to show that the distance between these dips in laminin fluorescence matched the distance between the micropores was performed. To do this a line was drawn through the laminin image in FIJI (ImageJ) and the plot profile function was used to get the mean fluorescent intensity at each point along the line. This data was then exported to Matlab where it was plotted inversely, so that the dips in fluorescence would be the peaks, and then the findpeaks function was used to identify the peaks. The distance between the peaks was averaged to find the average distance between the defects and that distance was compared to the distance between the micropores as measured by SEM imaging.

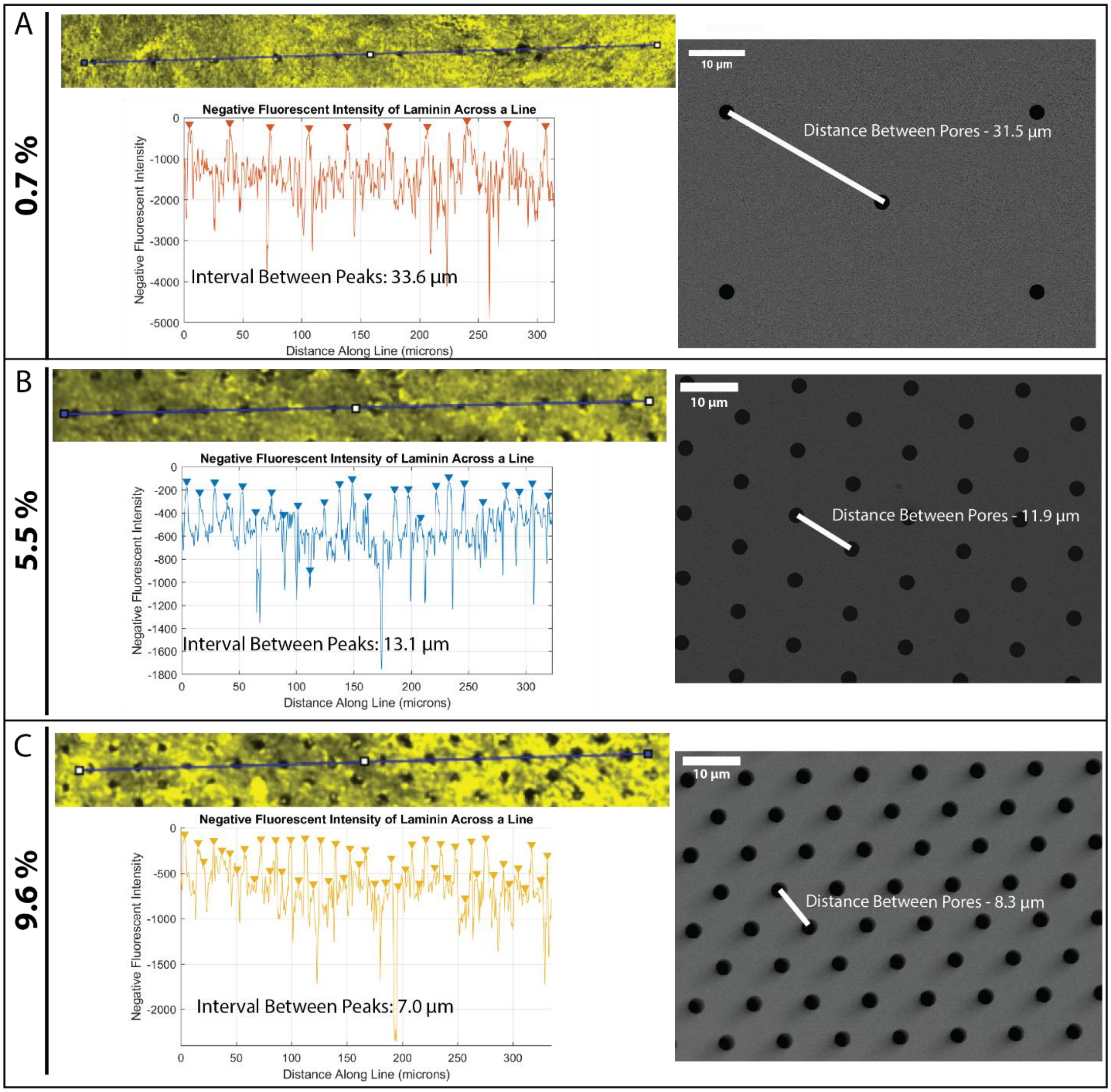


**Figure S1.** Validation of the distance between laminin defects on monoculture 3 µm DS membranes. Confocal imaging of laminin (yellow) with line drawn to show where the profile data is obtained. Graph of inverted mean fluorescent intensity profile so that peaks represent dips in fluorescence with peaks identified using Matlab's findpeaks function. The distance between these peaks was averaged to find the interval between peaks and is compared to the distance between the peaks as determined on SEM images. Analysis performed on 0.7% (A), 5.5 % (B), and 9.6% (C) 3 μm DS membranes.

Supplementary Section S2: Investigation of laminin expression by BPLCs alone.

To demonstrate that the fibrous laminin we see in the co-culture on dual scale membranes is a phenomenon at the interface of the coculture, and not merely a product of pericytes alone, we show representative images of laminin staining in BPLC only cultures and deep in the BPLC layer of the cocultures. The laminin expression is very similar in both of these settings, appearing limited to the cell body and more punctate than fibrous.

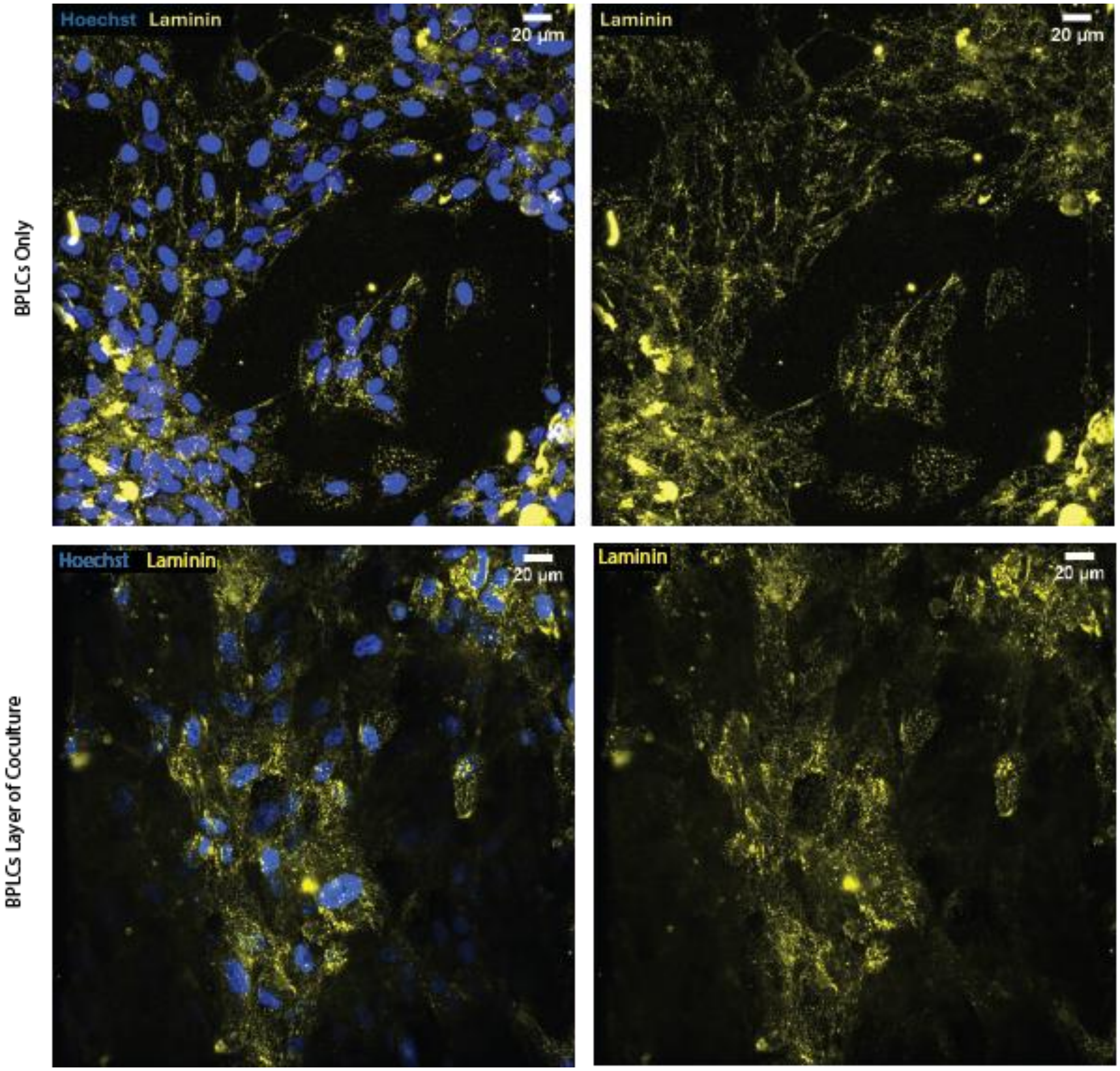


**Figure 2.** BPLCs alone produce different laminin texture to BMECs. Top: Confocal microscopy of laminin (yellow) staining of BPLC only culture. Bottom: Confocal microscopy of laminin staining of BPLC layer of coculture with BMECs. Both show punctate laminin expression around cells only, as opposed to more fibrous laminin seen near BMECs.
